# CRISPR-Cas9-mediated Large Cluster Deletion and Multiplex Genome Editing in *Paenibacillus polymyxa*

**DOI:** 10.1101/2021.08.06.455192

**Authors:** Meliawati Meliawati, Christa Teckentrup, Jochen Schmid

## Abstract

The use of molecular tools based on the clustered regularly interspaced short palindromic repeats-Cas (CRISPR-Cas) systems has rapidly advanced genetic engineering. These molecular biological tools have been applied for different genetic engineering purposes in multiple organisms, including the quite rarely explored *Paenibacillus polymyxa*. However, only limited studies on large cluster deletion and multiplex genome editing have been described for this highly interesting and versatile bacterium. Here, we demonstrate the utilization of a Cas9-based system to realize targeted deletions of four biosynthetic gene clusters in the range of 12 kb to 41 kb by the use of a single targeting sgRNA. Furthermore, we also harness the system for multiplex editing of genes and large genomic regions. Multiplex deletion was achieved with more than 80 % efficiency, while simultaneous integration at two distantly located sites was obtained with 58 % efficiency. The findings reported in this study are anticipated to accelerate future research in *P. polymyxa* and related species.

## Introduction

*Paenibacillus polymyxa* is a Gram-positive, facultative anaerobe, endospore forming, non-pathogenic soil bacterium. It is known for its nitrogen fixation capability which makes it attractive for utilization in modern agriculture. Furthermore, *P. polymyxa* naturally produces a wide range of valuable compounds such as exopolysaccharides (EPS), 2,3-butanediol, as well as antibiotics and antimicrobial peptides.^1^ *P. polymyxa* produces different types of EPS depending on the carbon source. These EPSs have unique characteristics and physiochemical properties which could be explored for different applications, such as cosmetics, coatings or within the agrochemical industry.^2^ As *P. polymyxa* gains more and more interests as biotechnological workhorse, the availability of stable and robust genetic tools is essential to further study and engineer the strain towards a versatile platform organism.^3^ While some molecular biology tools have been described for *Paenibacillus* species in the past,^4–6^ the efficiencies were quite low and further development is of highest interest.

The discovery of clustered regularly interspaced short palindromic repeats (CRISPR) has revolutionized biotechnology research in unprecedented ways. As an RNA-guided adaptive defense mechanism, CRISPR originates and naturally exists in many bacteria and archaea.^7^ Today, the basic mechanism has widely been adapted in different research fields and applied for genetic engineering of various organisms. Several CRISPR-associated (Cas) proteins have been described, but Cas9 is the most studied and well-characterized thus far. Cas9 belongs to the class 2 type II CRISPR-Cas system and has a single large effector protein.^8^ To realize a functional CRISPR-Cas9 system, the expression of Cas9 needs to be accompanied by guide RNA (gRNA) which consists of a CRISPR-RNA (crRNA) and trans-activating crRNA (tracrRNA) complex containing a spacer sequence homologous to the targeted region. A shorter and synthetic crRNA-tracrRNA, called single guide RNA (sgRNA), is commonly used today. The sgRNA acts as a guide for Cas9 to the targeted sequence. Cas9 itself is an endonuclease which recognizes the corresponding protospacer adjacent motive (PAM) sequence of 5′-NGG located downstream of the spacer. Upon binding to the target region, Cas9 is activated and generates a double strand DNA (dsDNA) break. This dsDNA break triggers the native DNA repair system which, depending on the organism, could be realized by homology-directed repair (HDR), non-homologous end joining, or alternative end-joining repair mechanisms.^9–11^ Further Cas proteins, such as Cas12a are also used in biotechnology research today and many variants such as nicking or deactivated Cas proteins exist.^12,13^ Thus, CRISPR-Cas systems can be used for genome editing, gene regulation as well as targeted labelling. This high versatility of CRISPR-Cas renders it a very powerful tool of modern biotechnology.

In 2017, the implementation of a CRISPR-based system to facilitate efficient genome editing in *P. polymyxa* was reported for the first time.^14^ The single-vector-based system was used to mediate single-gene deletions as well as the deletion of a genomic region of 18 kb by utilizing two targeting sgRNAs. Furthermore, a system for single and multiple gene regulation has been developed by employing the DNase-inactive variant of an optimized Cas12a (dCas12a) from *Acidaminococcus sp*.^13^ In addition, Kim *et al*.^15^ have recently developed a multiplex base editing tool for *P. polymyxa* by fusing the dCas9 with a cytidine deaminase to mediate C to T substitutions. However, no studies have reported multiplex gene deletions or integrations in *P. polymyxa* thus far. As modification of various genes is often needed for efficient metabolic engineering or the development of chassis organisms, the possibility to perform targeted multiple gene editing is highly attractive. In this study, we scope on the exploitation of multiplex genome editing to realize simultaneous gene deletions and integrations in *P. polymyxa*. The genes and cluster which are essential for the EPS production were selected as initial targets to allow for eased screening. In addition, we demonstrate the functionality of a single sgRNA-guided CRISPR-Cas9 approach to facilitate large cluster deletions, which will be important to realize mutant strains with minimized genomes towards the development of a robust and versatile chassis organism.

## Results and Discussion

Multiple genetic modifications are often required for microbial strain engineering. These modifications include, for example, gene deletions or integrations to direct the metabolism towards the desired pathway. Furthermore, deletion of non-essential genomic regions is of interest to generate minimalistic strains with superior performance.^16,17^ The availability of genetic tools which could facilitate reliable genome editing for such purposes is essential. While CRISPR-Cas-based systems have been used for genetic engineering of *P. polymyxa*, only limited studies reported their applications beyond the single-gene level. Here, we report the utilization of a Cas9-based system to realize deletion of large genomic regions as well as multiplex genome editing. The pCasPP plasmid was used as a base to realize the different genomic modifications.^14^ In brief, this plasmid contains the *cas9* gene from *Streptococcus pyogenes* which is codon-optimized for expression in *Streptomyces*. The expression of Cas9 is under the control of the constitutive *sgsE* promoter from *Geobacillus stearothermophilus* while the expression of the sgRNA is controlled by the constitutive *gapdh* promoter from *Eggerthella lenta*.^18,19^ Homologous regions of ∼1 kb upstream and downstream of the targeted regions were provided as HDR template. The spacer sequence was chosen 20-nt upstream of the NGG PAM sites.

In this study, we demonstrate the targeted deletions of four biosynthetic gene clusters (BGCs) which are responsible for the biosynthesis of paenan (*pep*), bacillibactin (*dhb*), fusaricidin (*fus*), and polymyxin (*pmx*) (Figure 1). To realize the cluster deletion, we employed the pCasPP-based plasmid only equipped with one targeting sgRNA. To analyze its efficiency, we first investigated the *pep* cluster which is 32.8 kb in size and comprises of 28 genes which support the production of paenan, the glucose-derived EPS produced by *P. polymyxa*. Deletion of the *pep* cluster will lead to elimination of paenan biosynthesis and therefore the mutants will not show a slimy phenotype when grown on EPS-inducing plates with glucose as the carbon source (EPS_glc_). Five different positions in the *pep* cluster were individually targeted to evaluate whether there is an effect of the location of the dsDNA break in the region on the editing efficiency. They were distributed within the cluster and had different distances to the homologous regions provided as repair template. The targeted sites were located in the *pepC* (*pep*-sg1), *pepH* (*pep*-sg2), *manC* (*pep*-sg3), *pepQ* (*pep*-sg4), and *pepT* (*pep*-sg5) genes, respectively (Table 1). The sgRNAs were chosen so that *pep*-sg1/sg5 and *pep*-sg2/sg4 had similar distances to the homologous regions, but located at the opposite ends of the cluster. Meanwhile, *pep*-sg3 was located in the middle of the cluster. Colonies in which the *pep* cluster is successfully deleted should result in a 3 kb PCR fragment when screened with primers binding outside of the homologous regions (Figure 2B). No amplicon should be obtained by the applied PCR reaction conditions for the wild type strain in which the *pep* cluster is still intact. Except for the targeting of *pep*-sg1, each conjugation round resulted in around 10 to 60 exconjugants with conjugation efficiencies between 1×10^−6^ to 4×10^−6^. Screening of the exconjugants showed that four out of five sgRNAs tested resulted in the deletion of the *pep* cluster. As expected, no slimy appearance was observed for the mutants on EPS_glc_ plate (Figure 2C). An editing efficiency of 90 % was obtained when *pep*-sg2 was targeted, while it was 100 % for *pep*-sg3, *pep*-sg4, and *pep*-sg5 (Figure 3A). Surprisingly, despite multiple trials, no exconjugants could be obtained when *pep*-sg1 was targeted. Thus, a tendency in which targeting the sites from the middle to 3′ end of *pep* cluster results in higher success rates to achieve the targeted deletion was observed. As *pep*-sg1 and *pep*-sg5 actually have similar distances to the homologous regions, the results suggest that distance of the cutting site to the homologous regions is not the only factor which determines the success of HDR.

**Table 1.** Overview of the biosynthetic gene clusters investigated in this study. Different positions in the clusters were targeted by individual sgRNA. The distance of each targeted site to the homologous regions are also listed.

| Gene cluster | Product | Size (kb) | Targeted site | Targeted gene | Predicted gene function | Distance of targeted site to homologous regions (in kb, 5' end; 3' end) |
| --- | --- | --- | --- | --- | --- | --- |
| ***pep*** | paenan | 32.8 | sg1 | *pepC* | Priming transferase | 2.4; 30.4 |
|  |  |  | sg2 | *pepH* | Flippase | 8.9; 23.9 |
|  |  |  | sg3 | *manC* | Mannose-1-phosphate guanylyltransferase | 16.4; 16.4 |
|  |  |  | sg4 | *pepQ* | Priming transferase | 24.0; 8.8 |
|  |  |  | sg5 | *pepT* | Transferase | 30.3; 2.5 |
| ***dhb*** | bacillibactin | 12.4 | sg1 | *dhbE* | 2,3-dihydroxybenzoate-AMP ligase | 3.2; 9.2 |
|  |  |  | sg2 | *dhbF* | Bacillibactin synthetase | 6.4; 6.0 |
| ***fus*** | fusaricidin | 30.7 | sg1 | *fusE* | Aldehyde dehydrogenase | 2.9; 27.8 |
|  |  |  | sg2 | *fusA* | Fusaricidin synthetase | 20.9; 9.8 |
| ***pmx*** | polymyxin | 41.0 | sg1 | *pmxA* | Polymyxin synthetase A | 2.4; 38.6 |
|  |  |  | sg2 | *pmxE* | Polymyxin synthetase E | 22.9; 18.1 |

**Figure 1.**
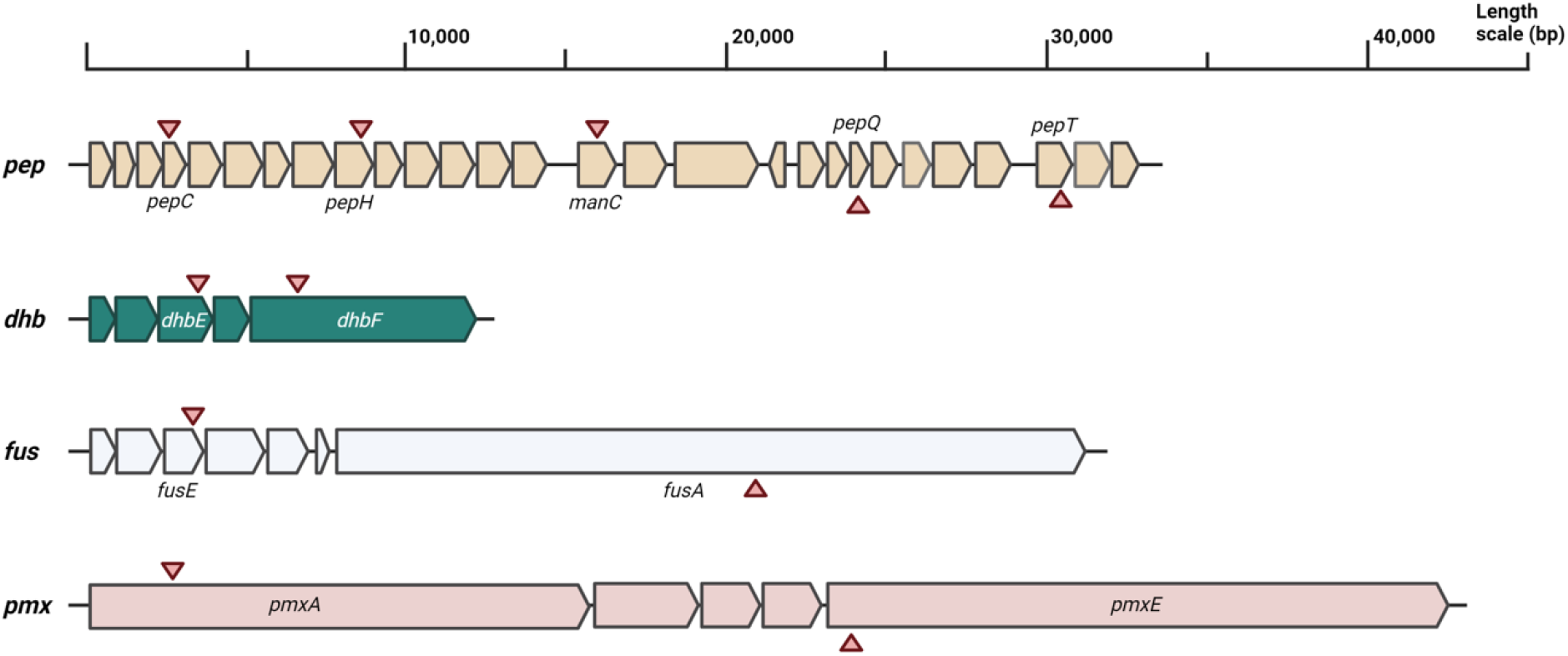
Schematic overview of the biosynthetic gene clusters of paenan (*pep*), bacillibactin (*dhb*), fusaricidin (*fus*), and polymyxin (*pmx*) of *P. polymyxa* DSM 365. The positions which were chosen as cutting sites within the respective genes are indicated by triangles.

**Figure 2.**
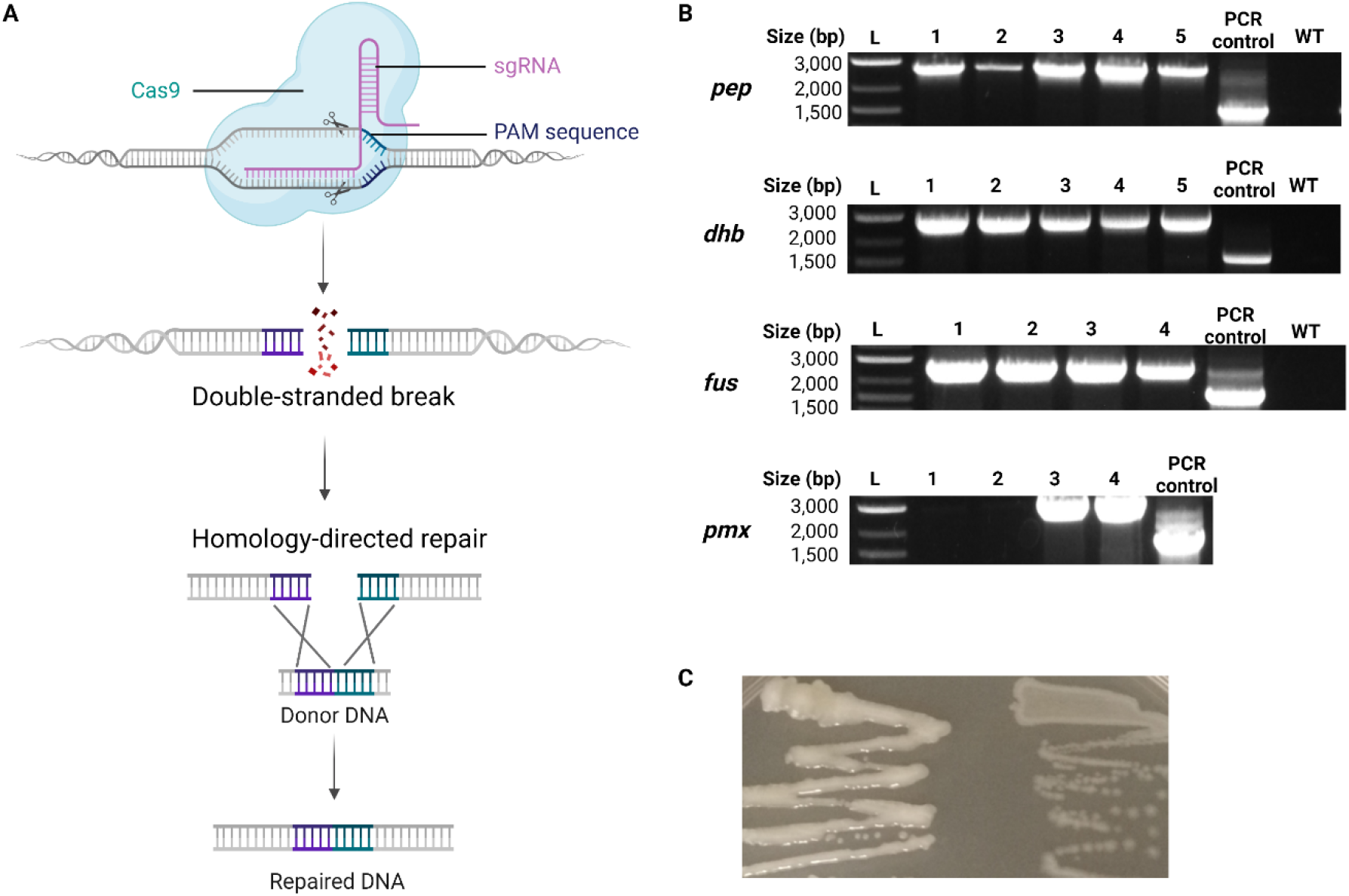
CRISPR-Cas9-mediated deletion of different biosynthetic gene clusters in *P. polymyxa*. (A) Schematic overview of Cas9-based targeted genome editing by employing the homology-directed repair mechanism. (B) Deletion of paenan (*pep*), fusaricidin (*fus*), bacillibactin (*dhb*), and polymyxin (*pmx*) clusters. PCR screening of randomly picked exconjugants (1-5), wild type (WT), and a PCR control. The PCR control was included to confirm the viability of the colony PCR from the WT strain by amplification of the 16S rRNA gene. The DNA ladder used was 1 kb Plus DNA Ladder from NEB. (C) Growth of WT (left) and mutant strains (right) on EPS-inducing plate with glucose as the carbon source.

**Figure 3.**
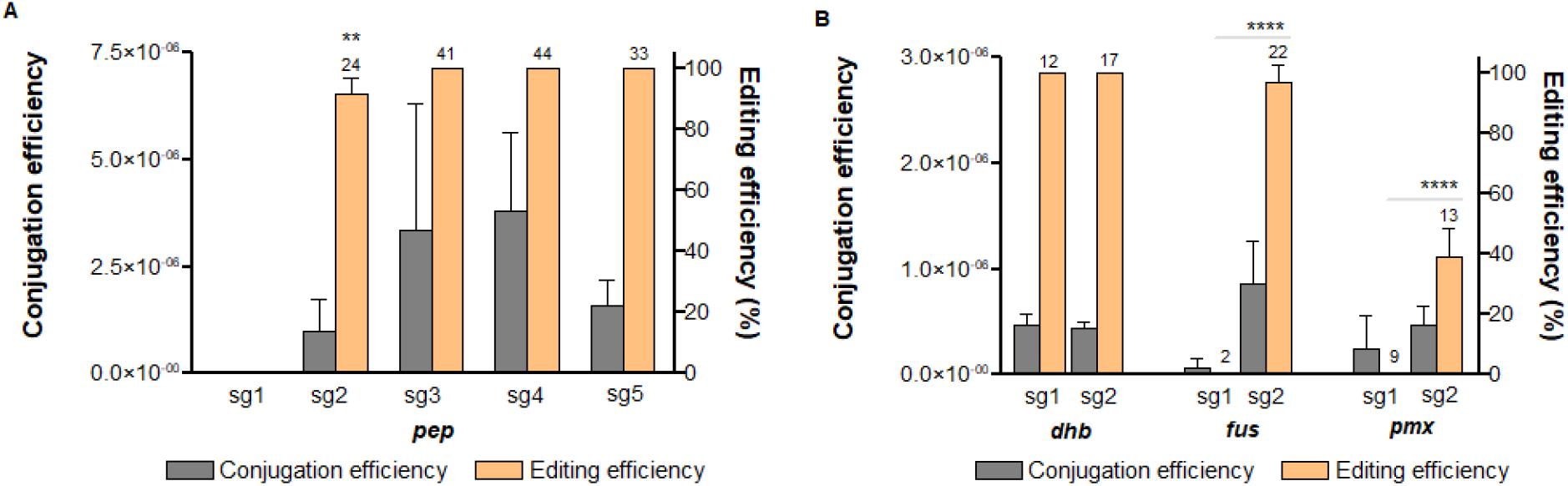
Targeted deletions of different biosynthetic gene clusters of *P. polymyxa*. Five different sites were individually targeted for the *pep* cluster (A) and two sites each for the *dhb, fus*, and *pmx* clusters (B). The conjugation efficiency (grey) and editing efficiency (orange) are provided on the Y-axes. The numbers on top of the editing efficiency bar indicate the total number of exconjugants screened. Error bars indicate the standard deviation. P value < 0.05 was considered in determining the significant differences between the editing efficiencies (** *p* < 0.01; **** *p* < 0.0001).

Subsequently, we targeted three other BGCs to see whether this phenomenon is also observed in other clusters or rather specific for the *pep* cluster. The clusters investigated were *dhb* (12 kb), *fus* (31 kb), and *pmx* (41 kb), which are responsible for the biosynthesis of bacillibactin, fusaricidin, and polymyxin, respectively. Two positions were individually targeted for each cluster, one located closer to the 5′ end (sg1) and one at the middle to 3′ end region (sg2) (Table 1). When targeting the *dhb* cluster, there was no difference in regards to the editing efficiency. Screening of the exconjugants showed that *dhb* deletion was achieved with 100 % efficiency when *dhb*-sg1 or *dhb*-sg2 was targeted. Interestingly, for *fus* and *pmx*, we observed the similar phenomenon as for the *pep* cluster, in which the deletion was not successful when *fus-*sg1 and *pmx*-sg1 were targeted. Meanwhile, it was possible to delete the cluster when targeting the sites located closer to the 3′ end (Figure 3B). In the case of *fus* cluster, the *fus*-sg2 is 20 kb and 10 kb away from the homologous regions, which is similar to *pep*-sg4 position targeted for the *pep* cluster deletion. By targeting *fus*-sg2, we successfully deleted the *fus* cluster with an average efficiency of 95 %. Meanwhile, deletion of *pmx* cluster was achieved with 39 % efficiency when targeting the *pmx*-sg2. Although deletion of the relatively small *dhb* cluster could be realized by targeting either *dhb*-sg1 or *dhb*-sg2, we observed that targeting larger clusters on the middle to 3′ end regions resulted in higher efficiencies. In fact, we were not able to achieve the deletion of *pep, fus*, and *pmx* clusters when targeting the sites located ∼2.5 kb away from the 5′ end (corresponding sg1s). Our findings suggest that the position of the cutting site has a significant impact on the success rate of large BGCs deletion by the use of a single sgRNA. We hypothesize that the three-dimensional structure of the BGCs might also be involved in this observation, but this requires further investigation. Based on these results, we thus recommend to target location in the middle to 3′ end regions when aiming for deletion of large BGCs, as it was proved to be successful for the deletion of four different BGCs investigated in this study.

Previous studies have demonstrated large cluster deletion in bacteria by the use of CRISPR-based systems. Commonly, dual targeting system with two sgRNAs are used in which the sgRNAs target the start and the end regions of the cluster, what allows a close proximity between the dsDNA break and the homologous regions.^12^ A study by So *et al*.^20^ showed that two sgRNAs were necessary to delete a 38 kb operon in *Bacillus subtilis* as utilization of single sgRNA failed in the targeted editing. Studies which describe the use of only one sgRNA-guided deletion of large clusters in bacteria are scarce. Recently, it was reported that a chromosomally integrated Cas12a was able to facilitate large cluster deletion in *B. subtilis* by the use of only one sgRNA.^21^ Here, we show highly efficient large cluster deletions with up to 100 % efficiencies by using a one plasmid system with only one sgRNA. We successfully demonstrate targeted deletions of BGCs up to 41 kb. The genome analysis of *P. polymyxa* by the use of antiSMASH^22^ and manual curation suggests that there exist at least 8 BGCs encoding for antimicrobial compounds with sizes ranging from 9 to 58 kb (Table S6). Therefore, the knowledge obtained in this study will substantially advance the application of Cas9-based approach for genome reduction in *P. polymyxa*, to create a robust production strain with a minimalistic genome.

The ability to perform simultaneous modifications would speed up and simplify the process in creating the strains of interest. Therefore, we were intrigued to exploit further our Cas9-based system for multiplexing purposes. Different strategies can be employed to realize the multiplexing, either by using single sgRNA expression cassettes or polycistronic sgRNAs. In the latter case, additional approach is needed to allow the processing of the gRNAs, for example by co-expression of Csy4 or fusing the gRNA with tRNA sequences.^23^ In this study, we applied the former strategy by using single sgRNA expression cassettes in a one plasmid system to allow simple cloning and transformation procedures (Figure 4A). First, we investigated the simultaneous deletion of *pepC* and *sacB* genes, which are essential for two different EPS biosynthesis pathways in *P. polymyxa*. As discussed above, *pepC* encodes for an initial glycosyltransferase required for paenan production. On the other hand, *sacB* encodes for a levansucrase which represents the single enzyme required for biosynthesis of levan, a sucrose-derived fructanosyl polymer.^24^ Disruption of these two genes will lead to a complete loss of EPS production in *P. polymyxa*.

**Figure 4.**
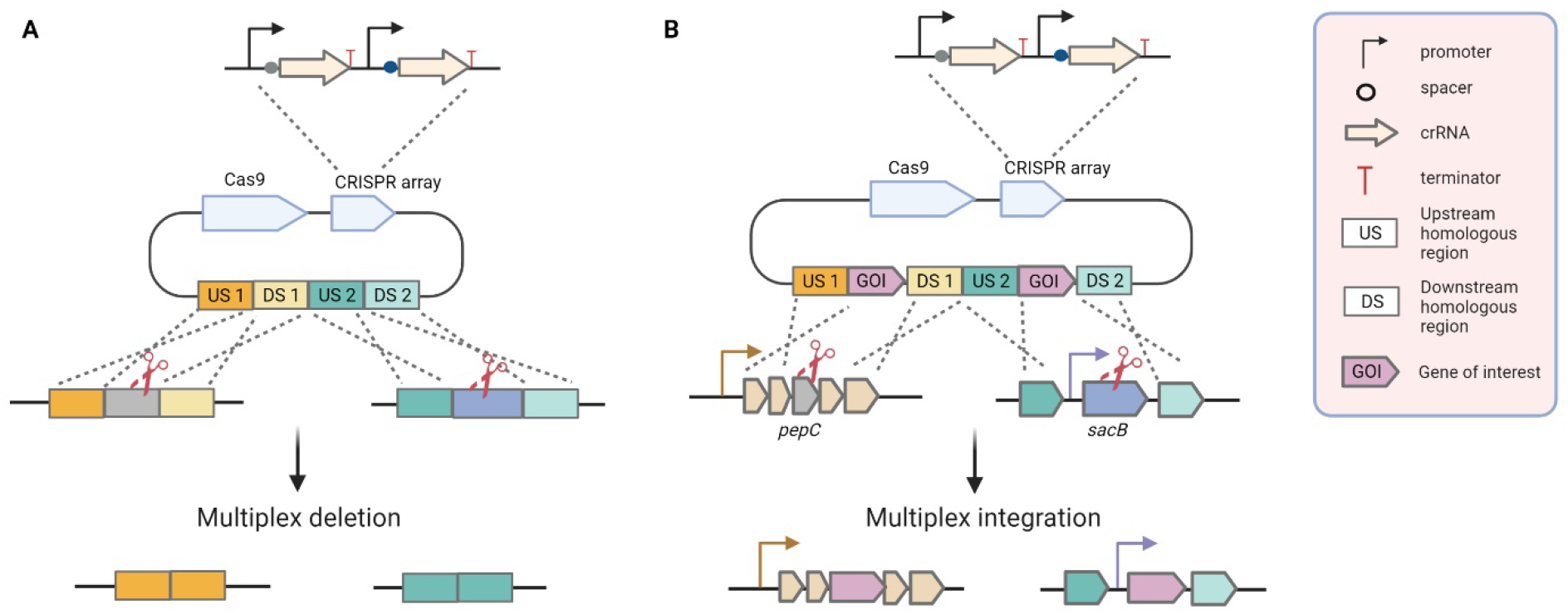
Strategy for Cas9-mediated multiplex genome editing by the use of one plasmid system to achieve two deletions (A) and two integrations (B). The plasmid harbors all elements needed for the targeted modifications including Cas9, CRISPR array, and homologous regions as repair template. The CRISPR array consists of two sgRNA expression cassettes, each of them containing a promoter, spacer sequence, crRNA, and a terminator.

Typically, conjugation efficiencies of around 1 ⨯ 10^−4^ are obtained for single-gene knockouts, although this number seems to vary for different target genes. Single-gene deletions of *pepC* and *sacB* were achieved with editing efficiencies of 94 % and 100 %, respectively. When aiming for simultaneous deletion of the two genes, the conjugation efficiency dropped significantly by about 70-fold compared to the single-gene deletions. As expected, deletion of *pepC* and *sacB* leads to a non-mucoid growth on EPS_glc_ and EPS with sucrose as the carbon source (EPS_suc_), what served as a first visual screening (Figure S1). Subsequently, the colonies were checked with PCR and sequencing to confirm the desired deletions. The results obtained from PCR and sequencing perfectly matched the visual plate-based screening, what underpins the reliability of the screening method. Despite the decrease in the conjugation efficiency, the editing efficiency of simultaneous gene deletions remained quite high with 85 % (Figure 5A).

**Figure 5.**
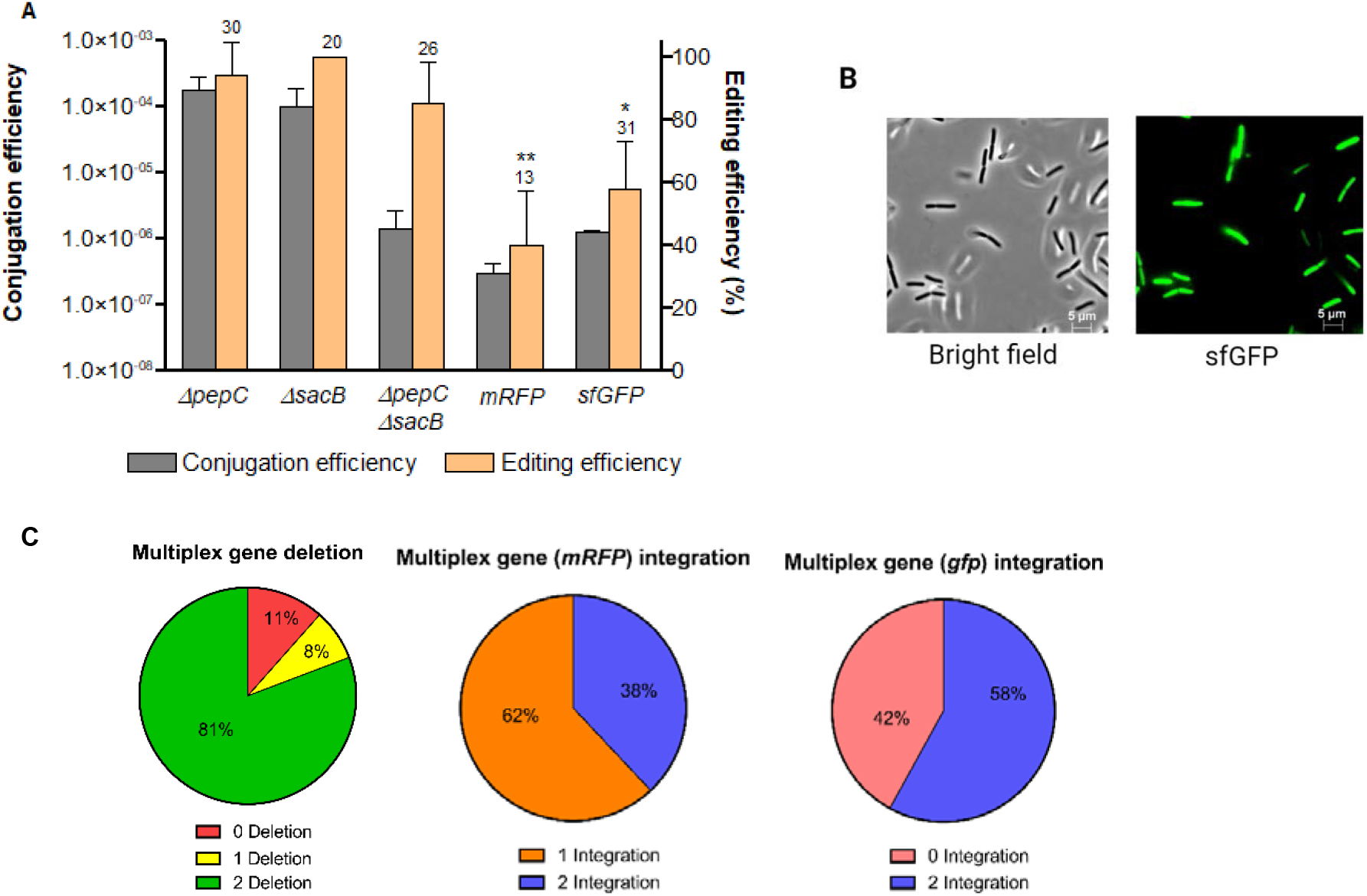
CRISPR-Cas9-based multiplex genome editing for *pepC* and *sacB* genes. (A) Conjugation and editing efficiencies of the single gene deletions, multiplex deletions, and multiplex integrations with *mRFP* or *sfGFP* as the gene of interest. The conjugation efficiency (grey) and editing efficiency (orange) are provided on the Y-axes. The numbers on top of the editing efficiency bar indicate the total number of exconjugants screened. Error bars indicate the standard deviation. P value < 0.05 was considered in determining the significant differences between the editing efficiencies (* *p* < 0.05; ** *p* < 0.01). Significant differences were obtained for multiplex *mRFP* integrations compared to the single gene deletions as well as multiplex deletions, also for multiplex *sfGFP* integrations compared to the single gene-deletions. (B) Microscopic images of *P. polymyxa* mutants with *sfGFP* replacing the *pepC* and *sacB* genes. (Left) Bright-field image; (right) GFP channel. (C) The proportion of wild type and mutants obtained from the screening process are provided as pie charts.

We further investigated whether the system could also be applied to realize multiplex gene integrations (Figure 4B). For this, *pepC* and *sacB* were targeted by using the same sgRNAs cassettes as used for the deletion experiments. In addition, the *mRFP* gene was placed in between the homologous regions of each targeted gene. Again, the conjugation efficiency decreased by 5-fold when attempting for multiplex gene integrations in comparison to the multiplex gene deletions. Nevertheless, simultaneous integrations of the *mRFP* was confirmed with an editing efficiency of 40 % (Figure 5A). Finally, in total 13 colonies were screened, for which 5 colonies showed the correct integration of the *mRFP* in the targeted positions of *pepC* and *sacB*. The other 8 colonies only had the *sacB* replaced by the *mRFP* gene while *pepC* was still intact at the original location. By sequencing the sgRNA cassette of the plasmid from the escapers, mutations in the sgRNA sequence for *pepC* were identified, which explained the single targeting of *sacB*. In these escapers, the sgRNA which was supposed to target the *pepC* was mutated to the sgRNA sequence targeting the *sacB*, which suggested that an unwanted recombination happened in the CRISPR array.

Despite the correct integration of the *mRFP* gene at desired positions in the genome as confirmed by sequencing (Figure S2), it was not functionally expressed. No fluorescence signal was detected in the *P. polymyxa* mutant, although red coloration was readily visible in the *Escherichia coli* strains during cloning and conjugation steps. Here, we integrated the *mRFP gene* in place of the *pepC* and *sacB* genes, and aimed to achieve its expression by employing the native expression system of *P. polymyxa*, what was not successful in this case. By that, we conducted additional experiments in which we integrated the *sfGFP* gene, by using similar approach as for the *mRFP* integration. Multiplex integration of the *sfGFP* by simultaneously replacing the *pepC* and *sacB* genes was achieved with an average editing efficiency of 58 % (Figure 5A). Moreover, the *sfGPP* was successfully expressed, as observed from the visualization by fluorescence microscopy (Figure 5B).

The differences in regard of expression of the chromosomally integrated *mRFP* and *sfGFP* genes will need further investigations, but can be linked to differences in the necessary promoter and RBS sequences. Furthermore, as the conjugation efficiency obtained from multiplexing approaches was rather low, optimization of transformation procedure might improve the efficiency for multiplex integration.^25,26^ Previous studies have reported CRISPR-Cas-based multiplex genetic engineering in different microorganisms.^27–29^ However, only a few demonstrated multiplex gene integration, which was mainly performed in yeast.^30,31^ While it has been reported in *E. coli*,^32^ to the best of our knowledge, our study is the first to utilize a Cas9-based system to achieve multiplex gene integrations in Gram-positive bacteria.

With the successful results of large cluster deletions and multiplexing, we were intrigued to see whether we could realize multiplexing beyond individual gene levels. Using the similar approach, we aimed to delete the *pep* and *dhb* clusters simultaneously. For this purpose, the respective positions of *pep*-sg4 and *dhb-*sg2 of the *pep* and *dhb* clusters were targeted, as described previously. The exconjugants obtained by this approach were then screened based on their mucoid/non-mucoid growth on EPS_glc_ plates, as well as by PCR and sequencing. Remarkably, simultaneous deletion of both clusters could be achieved with an editing efficiency of 83 % (Figure 6), which was comparable to the multiplex deletion of *pepC* and *sacB* genes. Among the 21 exconjugants screened, three escapers were identified for which one still had both clusters intact, while the other two had the *pep* cluster deleted but the *dhb* cluster intact. As observed for most of the escapers in this study, they were no longer able to grow on selection media containing neomycin, what in consequence hindered the investigation whether there was any mutation in the sgRNAs. We hypothesize that, in this case, mutations in the origin of replication on the Cas9-encoding plasmid might have occurred, as described by Uribe *et al*.^33^

**Figure 6.**
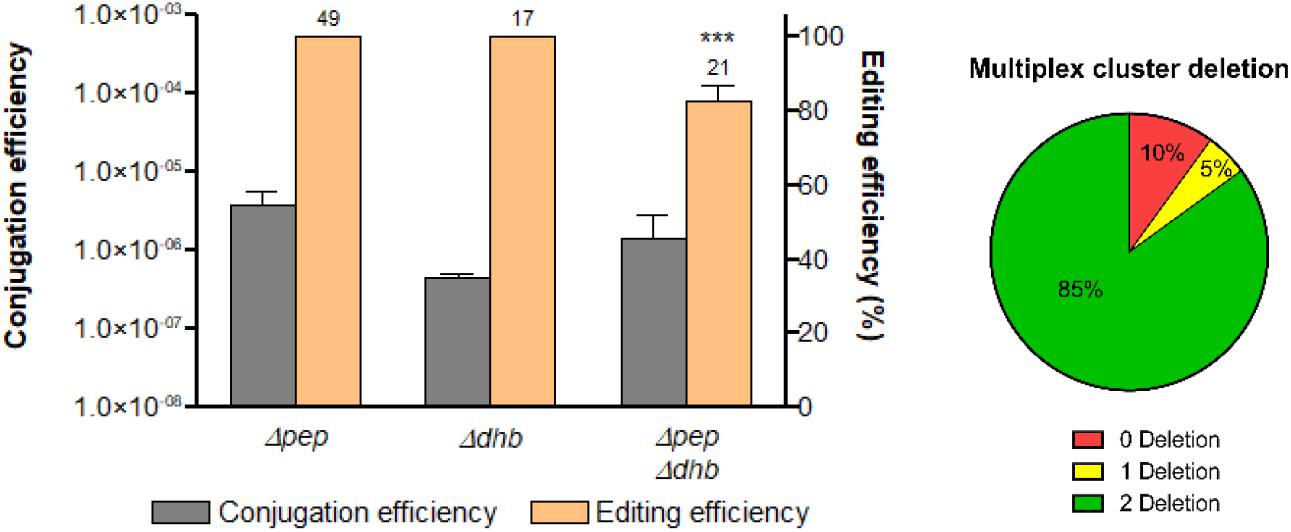
CRISPR-Cas9-based multiplex genome editing for *pep* and *dhb* clusters (B). The conjugation efficiency (grey) and editing efficiency (orange) are provided on the Y-axes. The numbers on top of the editing efficiency bar indicate the total number of exconjugants screened. Error bars indicate the standard deviation. P value < 0.05 was considered in determining the significant differences between the editing efficiencies (*** *p* < 0.001). The proportion of wild type and mutants obtained from the screening process are also provided as a pie chart.

To summarize, in this study, we demonstrate the utilization of our Cas9-based system for multiple purposes in *P. polymyxa*. In addition to single-gene deletions, we show that the system has high efficiencies for large cluster deletions as well as multiplex genome editing on gene and gene-cluster levels. Especially, we want to highlight the findings of the influence of the different targeting sgRNAs for different BGCs, in which it becomes obvious that targeting a region in the middle or closer to the 3′ end is more beneficial for large cluster deletion. The approach of multiplex cluster deletion allowed us to simultaneously delete two distantly located BGCs, which in total reduced 45 kb of the *P. polymyxa* genome. By that, we could show that this is a powerful approach for genome reduction towards specific minimalistic chassis variants. As a proof of principle, we also demonstrate multiplex gene integrations, which is rarely described in bacteria. Thus, the knowledge obtained in this study is expected to be substantial in strain development, especially when aiming for iterative genome editing. Multiplex integrations to simultaneously replace different BGCs would be an attractive direction to look into, but optimization of transformation efficiency is needed for this purpose. Finally, we anticipate a wider utilization of our Cas9-based system for future research in *P. polymyxa* and related species.

## Methods

### Strains and cultivation

*P. polymyxa* DSM 365 was obtained from the German Collection of Microorganisms and Cell Culture (DSMZ, Germany). *E. coli* Turbo (New England Biolabs) or *E. coli* Top10 (Invitrogen) was used for plasmid cloning and storage. *E. coli* S17-1 (ATCC 47055) was used to facilitate the conjugation of *P. polymyxa*. The strains were cultivated in LB media (10 g/L peptone, 5 g/L yeast extract, 5 g/L NaCl). For plate media, 1.5 % agar was added. If necessary, the media was supplemented with 50 µg/mL neomycin and/or 40 µg/mL polymyxin. *P. polymyxa* was cultivated at 30 °C and *E. coli* strains at 37 °C, unless stated otherwise. Liquid cultures were prepared in 3 mL media in 13 mL tubes and incubated at 250 rpm. To check for EPS formation, *P. polymyxa* was grown on EPS-inducing plates containing 30 g/L glucose or 30 g/L sucrose, 5 g/L peptone, 1.33 g/L MgSO_4_.7H_2_O, 0.05 g/L CaCl_2_, 1.67 g/L KH_2_PO_4_, 2 mL/L RPMI 1640 vitamin solution (Sigma-Aldrich), and 1 mL/L trace elements (2.5 g/L FeSO_4_.7H_2_O, 2.1 g/L C_4_H_4_Na_2_O_6_.2H_2_O, 1.8 g/L MnCl_2_.4H_2_O, 0.075 g/L CoCl_2_.6H_2_O, 0.031 g/L CuSO_4_.7H_2_O, 0.258 g/L H_3_BO_3_, 0.023 g/L Na_2_MoO_4_, 0.021 g/L ZnCl_2_).^14^ For longer storage, the strains were stored in 24 % glycerol and kept at -80 °C. All strains used in this study are listed in Table S1.

### Plasmid construction

All fragments for cloning purposes were amplified by the use of Accuzyme polymerase (Bioline). Gel purification was performed by using the Monarch Gel Purification Kit (NEB). Isolation of genomic DNA (gDNA) was performed with the DNeasy Blood & Tissue Kit (Qiagen). The pCasPP plasmid was used as vector for the different experiments in this study.^14^ The plasmid itself is 9.7 kb in size, containing Cas9 of *S. pyogenes* which was codon optimized for *Streptomyces* as well as sgRNA expression cassette.^18^ In addition, pCasPP also has the elements which support its replication in *E. coli* and *P. polymyxa* as well as origin of transfer for conjugation purpose. All the plasmids were assembled by isothermal assembly. Linear fragments were obtained from PCR amplifications using primers with suitable overlapping sequences. The spacer sequences were identified within the targeted genes and located directly upstream of the NGG PAM sites. In this study, 20-nt spacer sequences were used. Approximately 1 kb homologous regions upstream and downstream of the targeted regions were amplified from *P. polymyxa* gDNA and provided as repair template for the HDR. For multiplex integrations experiment, the *mRFP* was amplified from the pLux01 plasmid, which was a gift from Tom Ellis (Addgene plasmid #78281),^34^ and placed in between the homologous regions. Finally, all fragments were mixed together in the isothermal assembly master mix and incubated at 50 °C for 1 hour followed by transformation of *E. coli* Turbo or *E. coli* Top10 via heat shock method. Transformed cells were plated on LB agar containing 50 µg/mL of neomycin. The transformants were initially screened with colony PCR using GoTaq polymerase (Promega) then plasmid isolation was performed by using GeneJET Plasmid Miniprep Kit (Thermo Fisher Scientific). Subsequently, the plasmid was checked by sequencing to confirm that the correct plasmid was assembled. Upon confirmation, *E. coli* S17-1 was transformed with the plasmid, also by using heat shock method. All primers synthesis and sequencing analysis were performed by Microsynth AG (Switzerland). In silico primer design, cloning, and sequence alignment were performed by using SnapGene version 5.1.5. The spacer sequences were selected based on their respective positions in the targeted regions as well as high on-target scores as analyzed by Benchling platform. The figures were created by using BioRender. The plasmids, oligonucleotides, and spacer sequences used in this study are listed in Table S1-S3. Detailed information of the materials used in this study is provided in Table S4.

### Conjugation

*P. polymyxa* transformation was performed via conjugation by the use of *E. coli* S17-1 as donor strain. Single colonies of the strains were inoculated in 3 mL LB media, with or without 50 µg/mL of neomycin, and cultivated overnight. The overnight culture was diluted 1:100 in fresh media then cultivated for 4 h at 37 °C and 250 rpm. Subsequently, 900 µL of *P. polymyxa* culture was incubated at 42 °C for 15 min then mixed with 300 µl of the *E. coli* S17-1 harboring the plasmid of interest. The mixture was centrifuged at 8,000 rpm for 3 min then the pellet was resuspended in 100 µL of LB media. The solution was dropped on an LB plate and incubated overnight at 30 °C. Afterwards, the culture was scrapped off the plate and resuspended in 150 µL of LB media, Subsequently, the culture was plated on LB plates containing 50 µg/mL of neomycin and 40 µg/mL of polymyxin, followed by incubation for 48 h at 30 °C. If necessary, plating with serial dilutions was performed to obtain countable colonies on the plates. All experiments were performed in three biological replicates. Conjugation efficiency was calculated as the total number of exconjugants per viable recipient cells. Editing efficiency was calculated from the number of correct mutants divided by total number of exconjugants screened from each conjugation round. The numbers of exconjugants as well as conjugation and editing efficiencies obtained from the different experiments are provided in Table S5.

### Screening of *P. polymyxa* exconjugants

Screening of the exconjugants was mainly performed by PCR and sequencing. In addition, the plate-based assays were also used to screen for mutants whose EPS-related genes or cluster were deleted. For this, the colonies were streaked on EPS-inducing plates to check for the loss of slimy phenotype which indicated the elimination of EPS formation. An EPS_glc_ plate was used for initial screening of the *pepC* gene or *pep* cluster knockout. In addition, an EPS_suc_ plate was used to screen for the Δ*sacB* Δ*pepC* double knockout mutants. The colonies were incubated at 30 °C overnight. Subsequently, the colonies were checked via colony PCR to verify the accuracy of the plate-based assay. Colony PCR was performed using primers which bind outside of the homologous regions provided as repair template. The resulting fragment was purified from the gel and sent for sequencing to confirm the modifications. Finally, gDNA was isolated from some of the colonies and again checked with PCR and sequencing to reconfirm the colony PCR results. Analysis with fluorescence microscopy was performed by using Carl Zeiss HBO 100 microscope. Microscopic images were processed with AxioVision imaging software.

## Statistical analysis

Statistical analysis was performed using GraphPad Prism version 7. Editing efficiencies of different experiments with the same scope were compared using One-Way ANOVA with Tukey correction. P value < 0.05 was considered significant.

## Associated Content

Supporting Information

## Author Information

### Author Contributions

MM and JS designed the experiments and wrote the manuscript. MM and CT performed the experiments and analyzed the data. All authors read and approved the final manuscript.

### Notes

The authors declare no competing financial interests.

## Acknowledgements

This study is part of the German Federal Ministry of Education and Research (BMBF) funded project Polymore with the number 031B0855A.

