## Supplementary material for "CRISPR-Cas9-mediated Large Cluster Deletion and Multiplex Genome Editing in *Paenibacillus polymyxa*": BioRxiv_Supporting information.pdf

**Table S1**. List of bacterial strains and plasmids.

| **Bacterial strains** | **Genotype** | **Purpose** | **Reference** |
|---|---|---|---|
| *E. coli* Turbo (NEB) | F' proA+B+ lacIq ΔlacZM15 / fhuA2 Δ(lac-proAB) glnV galK16 galE15 R(zgb-210::Tn10)TetS endA1 thi-1 Δ(hsdS-mcrB)5 | Cloning host | NEB |
| *E. coli* Top10 (Invitrogen) | F-*mcrA* Δ(*mrr-hsd*RMS-*mcr*BC) Φ80*Lac*ZΔM15 Δ *Lac*X74 *rec*A1 *ara*D139 Δ(*araleu*) 7697 *gal*U *gal*K *rps*L (StrR) *end*A1 *nup*G | Cloning host | Invitrogen |
| *E. coli* S17-1 (ATCC 47055) | recA pro hsdR RP42Tc::Mu-Km::Tn7 integrated into the chromosome | Conjugation strain | ATCC |
| *P. polymyxa* DSM 365 | Wild type | Final host | DSMZ |
| **Plasmids** | **Size (bp)** | **Purpose** | **Reference** |
| pCasPP | 9,663 | Plasmid vector for Cas9-based genome editing in *P. polymyxa* | [1] |
| pLux01 | 4,806 | Source of *mRFP* gene | [2] |
| pCasPP-*pepC* | 11,125 | Deletion of *pepC* gene | [1] |
| pCasPP-*sacB* | 11,211 | Deletion of *sacB* gene | [3] |
| pCasPP-*pep*sg1 | 11,655 | Deletion of *pep* cluster, targeting *pep*-sg1 position | This study |
| pCasPP-*pep*sg2 | 11,655 | Deletion of *pep* cluster, targeting *pep*-sg2 position | This study |
| pCasPP-*pep*sg3 | 11,655 | Deletion of *pep* cluster, targeting *pep*-sg3 position | This study |
| pCasPP-*pep*sg4 | 11,655 | Deletion of *pep* cluster, targeting *pep*-sg4 position | This study |
| pCasPP-*pep*sg5 | 11,655 | Deletion of *pep* cluster, targeting *pep*-sg5 position | This study |
| pCasPP-*dhb*sg1 | 11,498 | Deletion of *dhb* cluster, targeting *dhb*-sg1 position | This study |
| pCasPP-*dhb*sg2 | 11,498 | Deletion of *dhb* cluster, targeting *dhb*-sg2 position | This study |
| pCasPP-*fus*sg1 | 11,414 | Deletion of *fus* cluster, targeting *fus*-sg1 position | This study |
| pCasPP-*fus*sg2 | 11,414 | Deletion of *fus* cluster, targeting *fus*-sg2 position | This study |
| pCasPP-*pmx*sg1 | 11,631 | Deletion of *pmx* cluster, targeting *pmx*-sg1 position | This study |
| pCasPP-*pmx*sg2 | 11,631 | Deletion of *pmx* cluster, targeting *pmx*-sg2 position | This study |
| pCasPP-*pepC-sacB* | 13,533 | Multiplex deletion of *pepC* and *sacB* gene | This study |
| pCasPP-*pepC::mRFP sacB::mRFP* | 14,980 | Multiplex integration of *mRFP* to replace *pepC* and *sacB* genes | This study |
| pCasPP-*pepC::sfGFP sacB::sfGFP* | 15,058 | Multiplex integration of *sfGFP* to replace *pepC* and *sacB* genes | This study |
| pCasPP-*pep-dhb* | 14,358 | Multiplex deletion of *pep* and *dhb* clusters | This study |

**Table S2**. List of oligonucleotides.

| Plasmid | Primers | | |
|---|---|---|---|
| | Name | Sequences (5′→ 3′) | Purpose |
| pCasPP-*pep*sg1 | pCasPP_bb2_fw | CTGTTACAGGCATATTCATATCAATGTCG | plasmid construction |
| | pCasPP_bb1_rev | CCTCACCTCCTGCTACCATTCCTAGTCGGCGGGCTTGATGCG | |
| | pCasPP_bb3_rev | CGACATTGATATGAATATGCCTGTAACAG | |
| | pCasPP_bb2_fw | TCTAGTGGCCAGGAACCGTAAAAAGG | |
| | clu2_DS_fw | AGTCGGTAAAAGCAGGATGC | |
| | ugdh2_rev | TACGGTTCCTGGCCACTAGATTACGGATTTACTACAGACTGATG | |
| | clu1_US_rev | GCATCCTGCTTTTACCGACTTGTTGATTATGGAAAAAGAAATCGAAAAAAGCC | |
| | pepA_US_fw | GAATGGTAGCAGGAGGTGAGG | |
| pCasPP-*pep*sg2 | pCasPP_bb2_fw | CTGTTACAGGCATATTCATATCAATGTCG | plasmid construction |
| | pepH_sgRNA | CGAAGACAGCATTGCTATCCGCGTATCCCCTTTCAGATACTCG | |
| | pepH_sgRNA_fw | GGATAGCAATGCTGTCTTCGGTTTTAGAGCTAGAAATAGCAAGTTAAAATAAGGC | |
| | pCasPP_bb3_rev | CGACATTGATATGAATATGCCTGTAACAG | |
| pCasPP-*pep*sg3 | pCasPP_bb2_fw | CTGTTACAGGCATATTCATATCAATGTCG | plasmid construction |
| | manC_sgRNA | TCGTCATATTTGACATAATCGCGTATCCCCTTTCAGATACTCG | |
| | manC_sgRNA_fw | GATTATGTCAAATATGACGAGTTTTAGAGCTAGAAATAGCAAGTTAAAATAAGGC | |
| | pCasPP_bb3_rev | CGACATTGATATGAATATGCCTGTAACAG | |
| pCasPP-*pep*sg4 | pCasPP_bb2_fw | CTGTTACAGGCATATTCATATCAATGTCG | plasmid construction |
| | pCasPP_bb1_rev | CCTCACCTCCTGCTACCATTCCTAGTCGGCGGGCTTGATGCG | |
| | pCasPP_bb3_rev | CGACATTGATATGAATATGCCTGTAACAG | |
| | pepA_US_fw | GAATGGTAGCAGGAGGTGAGG | |
| pCasPP-*pep*sg5 | pCasPP_bb2_fw | CTGTTACAGGCATATTCATATCAATGTCG | plasmid construction |
| | pepT_sgRNA_rev | AGCTGCCTTTTGTTAATGGTGCGTATCCCCTTTCAGATACTCG | |
| | pepT_sgRNA_fw | ACCATTAACAAAAGGCAGCTGTTTTAGAGCTAGAAATAGCAAGTTAAAATAAGGC | |
| | pCasPP_bb3_rev | CGACATTGATATGAATATGCCTGTAACAG | |
| | clu_KO_fw | ATCCGTCATGGATTGGCCAAG | screening of *pep* cluster deletion |
| | clu_KO_rev | CGGATGATACATCGCATTCG | |
| pCasPP-*dhb*sg1 | dhb_sgRNA_fw | GGAATGTGACAACGACATGGGTTTTAGAGCTAGAAATAGCAAGTTAAAATAAGGC | plasmid construction |
| | oriT_rev | CTAGtCGGCGGGCTTGATGC | |
| | pCasPP_bb2_fw | CTGTTACAGGCATATTCATATCAATGTCG | |
| | dhb_sgRNA_rev | CCATGTCGTTGTCACATTCCGCGTATCCCCTTTCAGATACTC | |
| | pCasPP_bb2_fw | TCTAGTGGCCAGGAACCGTAAAAAGG | |
| | pCasPP_bb3_rev | CGACATTGATATGAATATGCCTGTAACAG | |
| | dhb_DS_fw | CTGGAAAGGGTGGAATGAACATGATTAATCCTTTTGAGCAGAATGAAGG | |
| | dhb_DS_rev | TACGGTTCCTGGCCACTAGATTGCTCAATAGATGCATTACGAC | |
| | dhb_US_fw | TTCATCGTTCCAGTGATGCAC | |
| | dhb_US_rev | GTTCATTCCACCCTTTCCAGC | |
| pCasPP-*dhb*sg2 | pCasPP_bb2_fw | CTGTTACAGGCATATTCATATCAATGTCG | plasmid construction |
| | dhb_sg2_rev | CTTCTTTGTCCTCATCATCCGCGTATCCCCTTTCAGATACTC | |

| | | | |
|---|---|---|---|
| | dhb_sg2_fw | GGATGATGAGGACAAAGAAGGTTTTAGAGCTAGAAATAGCAAGTTAAAATAAGGC | |
| | pCasPP_bb3_rev | CGACATTGATATGAATATGCCTGTAACAG | |
| | dhb_KO_check_fw | GAAGATTGGTTGGCCTTCATC | screening of *dhb* cluster deletion |
| | dhb_KO_check_rev | CAAGGCCGTGTCAAATCGTAG | |
| pCasPP-*fus*sg1 | fus_sg1_fw | CACCGGGAGCTATACCAATGGTTTTAGAGCTAGAAATAGCAAGTTAAAATAAGGC | plasmid construction |
| | oriT_rev | CTAGTCGGCGGGCTTGATGC | |
| | pCasPP_bb2_fw | CTGTTACAGGCATATTCATATCAATGTCG | |
| | fus_sg1_rev | CATTGGTATAGCTCCCGGTGGCGTATCCCCTTTCAGATACTCG | |
| | pCasPP_bb2_fw | TCTAGTGGCCAGGAACCGTAAAAAGG | |
| | pCasPP_bb3_rev | CGACATTGATATGAATATGCCTGTAACAG | |
| | fus_DS_fw | GATGACAGACAGCATACATGCTGGTGTCGGTGATGAGTTC | |
| | fus_DS_rev | TACGGTTCCTGGCCACTAGAGGAGCTAGCCACAAACCATG | |
| | fus_US_fw | GCATCAAGCCCGCCGACTAGATGTACCCGTGCTGACTAGC | |
| | fus_US_rev | CATGTATGCTGTCTGTCATCTCAC | |
| pCasPP-*fus*sg2 | pCasPP_bb2_fw | CTGTTACAGGCATATTCATATCAATGTCG | plasmid construction |
| | fus_sg2_rev | TCATGCTCTATCGGAAAGACGCGTATCCCCTTTCAGATACTCG | |
| | fus_sg2_fw | GTCTTTCCGATAGAGCATGAGTTTTAGAGCTAGAAATAGCAAGTTAAAATAAGGC | |
| | pCasPP_bb3_rev | CGACATTGATATGAATATGCCTGTAACAG | |
| | fus_check_fw | TATGCCATTCGTCCACGCTC | screening of *fus* cluster deletion |
| | fus_check_rev | CATCTAAGAACGCCGTGAGATC | |
| pCasPP-*pmx*sg1 | pmx_sg1_fw | GATCATTTCTTCGAGCTGGGGTTTTAGAGCTAGAAATAGCAAGTTAAAATAAGGC | plasmid construction |
| | oriT_rev | CTAGTCGGCGGGCTTGATGC | |
| | pCasPP_bb2_fw | CTGTTACAGGCATATTCATATCAATGTCG | |
| | pmx_sg1_rev | CCCAGCTCGAAGAAATGATCGCGTATCCCCTTTCAGATACTCG | |
| | pCasPP_bb2_fw | TCTAGTGGCCAGGAACCGTAAAAAGG | |
| | pCasPP_bb3_rev | CGACATTGATATGAATATGCCTGTAACAG | |
| | pmx_DS_fw | ATTGACATTGTTGTCACATGCAAGAAGTAATGGCCGATCGTTG | |
| | pmx_DS_rev | TACGGTTCCTGGCCACTAGACATCGTACCAAGGTCCATCTC | |
| | pmx_US_fw | GCATCAAGCCCGCCGACTAGCCAAGTGGGTACTGGATATGACTC | |
| | pmx_US_rev | CATGTGACAACAATGTCAATGGC | |
| pCasPP-*pmx*sg2 | pCasPP_bb2_fw | CTGTTACAGGCATATTCATATCAATGTCG | plasmid construction |
| | poly_sg1_rev | CCATGTACGTTTTATTGTACGCGTATCCCCTTTCAGATACTCG | |
| | poly_sg1_fw | GTACAATAAAACGTACATGGGTTTTAGAGCTAGAAATAGCAAGTTAAAATAAGGC | |
| | pCasPP_bb3_rev | CGACATTGATATGAATATGCCTGTAACAG | |
| | pmx_check_fw | GATAGAGTGGGCACTTATTACGG | screening of *pmx* cluster deletion |
| | pmx_check_rev | CTTCTCTAGTGATCCCCTTACTCG | |
| pCasPP-*pepC-sacB* | pCasPP_bb2_fw | CTGTTACAGGCATATTCATATCAATGTCG | plasmid construction |
| | sacB_ter_rev | TAAAAAACGCCCGGCGGCAAC | |
| | gapdh_prom_fw | TTGCCGCCGGGCGTTTTTTAGCTGCTCCTTCGGTCGGAC | |
| | pepC_flanks_rev | TCCATACAACAATGTAGACATCACCCCGCAGCTGACTCG | |
| | pCasPP_bb3_rev | CGACATTGATATGAATATGCCTGTAACAG | |
| | sacB_US_fw | TGTCTACATTGTTGTATGGACTGGG | |
| | pCasPP_bb2_fw | CTGTTACAGGCATATTCATATCAATGTCG | plasmid construction |

| | | | |
|---|---|---|---|
| pCasPP-*pepC::mRFP-sacB::mRFP* | pepC_sg1_R_AA1 | AAACCCGGTAAATTTTCCATGCTC | |
| | pCasPP_bb3_rev | CGACATTGATATGAATATGCCTGTAACAG | |
| | sacB_DS_mRFP_fw | CGTCACTCCACCGGTGCTTAAGTAATGCAGTATGTCCGAAGG | |
| | mRFP_prom_fw | TTTATATACTAGAGACCTGTAGGATCG | |
| | mRFP1_REV | TTAAGCACCGGTGGAGTGACG | |
| | pepC_DS_mRFP_fw | GTCACTCCACCGGTGCTTAAGGATGCTTATTGATGGATGCATTC | |
| | sacB_US_mRFP_rev | TACAGGTCTCTAGTATATAAACGATTGATATAGACTGAAAAGCCC | |
| | pepC_sg1_F_AA1 | ACGCGAGCATGGAAAATTTACCGG | |
| | iGT1_KOproof_F | GCAACTCATGGAGCAGGCCAGTGCGACG | screening of *pepC* gene deletion |
| | iGT1_KOproof_R | GCTGTATGGTGTTTATTCTAGTAGTCCAAGG | |
| | sacB_KOproof_F | AATTATCGCATTGCTGCCCAGACAG | screening of *sacB* gene deletion |
| | sacB_KOproof_R | AGATCGGGTTGCTACCAATCTACCG | |
| pCasPP-*pepC::sfGFP-sacB::sfGFP* | pCasPP_bb2_fw | CTGTTACAGGCATATTCATATCAATGTCG | |
| | pCasPP_bb3_rev | CGACATTGATATGAATATGCCTGTAACAG | plasmid construction |
| | pk19_bb1_fw | CCTACTTCACCTATCCTGCCC | |
| | pk19_bb2_rev | GGGCAGGATAGGTGAAGTAGG | |
| | GFP_fw | ATGCGTAAAGGCGAAGAGCTG | |
| | GFP_rev | TCATTTGTACAGTTCATCCATACCATGC | |
| | sacB_US_fw | TGTCTACATTGTTGTATGGACTGGG | |
| | pepC_flanks_rev | TCCATACAACAATGTAGACATCACCCCGCAGCTGACTCG | |
| | sacB_DS_rev | TACGGTTCCTGGCCACTAGAGCCACTCAGGTTCACTGTGATC | |
| | pCasPP_bb2_fw | TCTAGTGGCCAGGAACCGTAAAAAGG | |
| | pCasPP_bb3_rev | CGACATTGATATGAATATGCCTGTAACAG | |
| pCasPP-*pep-dhb* | pCasPP_bb2_fw | CTGTTACAGGCATATTCATATCAATGTCG | plasmid construction |
| | ter_rev | TAAAAAACGCCCGGCGGCAAC | |
| | gapdh_prom_fw | TTGCCGCCGGGCGTTTTTTAGCTGCTCCTTCGGTCGGAC | |
| | dhb_clu1_DS_rev | CTCACCTCCTGCTACCATTCTTGCTCAATAGATGCATTACGAC | |
| | pCasPP_bb3_rev | CGACATTGATATGAATATGCCTGTAACAG | |
| | pepA_US_fw | GAATGGTAGCAGGAGGTGAGG | |
| | 27f | AGAGTTTGATCMTGGCTCAG | amplification of 16S rRNA |
| | 1492r | GGTTACCTTGTTACGACTT | |

**Table S3**. List of spacer sequences for Cas9-mediated targeted modifications.

| Targeted gene/cluster | Targeted position | Spacer sequence (5′ → 3′) | PAM site | On-target score* |
|---|---|---|---|---|
| *pep* cluster | *pep*-sg1 | GAGCATGGAAAATTTACCGG | AGG | 78.0 |
| | *pep*-sg2 | GGATAGCAATGCTGTCTTCG | GGG | 63.6 |
| | *pep*-sg3 | GATTATGTCAAATATGACGA | GGG | 68.7 |
| | *pep*-sg4 | CGACCCACACGGGTAATTCG | GGG | 69.5 |
| | *pep*-sg5 | ACCATTAACAAAAGGCAGCT | AGG | 61.5 |
| *dhb* cluster | *dhb*-sg1 | GGATGATGAGGACAAAGAAG | TGG | 60.7 |
| | *dhb*-sg2 | GGAATGTGACAACGACATGG | TGG | 77.2 |
| *fus* cluster | *fus*-sg1 | CACCGGGAGCTATACCAATG | CGG | 72.2 |
| | *fus*-sg2 | GTCTTTCCGATAGAGCATGA | CGG | 69.3 |
| *pmx* cluster | *pmx*-sg1 | CACCGGGAGCTATACCAATG | CGG | 71.6 |
| | *pmx*-sg2 | GTACAATAAAACGTACATGG | CGG | 79.5 |
| *pepC* gene | | GAGCATGGAAAATTTACCGG | AGG | 78.0 |
| *sacB* gene | | ATTGTAGACGCATCGAAGGA | CGG | 60.7 |

*Analyzed using Benchling platform.

**Table S4**. List of materials used in this study.

| Type | Materials | Catalog number | Supplier |
|---|---|---|---|
| Chemicals | Tryptone/Peptone | 8952 | Carl Roth |
| | Yeast extract | 2363 | Carl Roth |
| | Sodium chloride | 3957 | Carl Roth |
| | Neomycin | 2363 | Carl Roth |
| | Polymyxin | 0235 | Carl Roth |
| | Glucose | 6780 | Carl Roth |
| | Sucrose | 4621 | Carl Roth |
| | Magnesium sulfate heptahydrate | PO27.2 | Carl Roth |
| | Calcium chloride dihydrate | CN93 | Carl Roth |
| | Potassium dihydrogen phosphate | 3904 | Carl Roth |
| | Vitamin solution RPMI 1640 | R7256 | Sigma-Aldrich |
| | Iron (II) sulfate heptahydrate | 12354 | Fluka |
| | Sodium molybdate dihydrate | M1651 | Sigma-Aldrich |
| | Manganese (II) chloride tetrahydrate | T881.1 | Carl Roth |
| | Cobalt (II) chloride hexahydrate | 12914 | Fluka |
| | Copper (II) sulfate pentahydrate | 2790 | Merck |
| | Boric acid | 6943 | Carl Roth |
| | Potassium sodium tartrate tetrahydrate | 108087 | Merck |
| | Zinc chloride | B722516 | Merck |
| | Glycerol | 7530 | Carl Roth |
| Enzymes and kits | Accuzyme DNA polymerase | BIO-21052 | Bioline |
| | GoTaq DNA polymerase | M3005 | Promega |
| | GeneJet Plasmid Miniprep | K0503 | Thermo Fisher Scientific |
| | Monarch DNA Gel Extraction | T1020L | New England Biolabs |
| | DNeasy Blood & Tissue Kit | 69504 | Qiagen |

**Table S5**. List of conjugation and editing efficiencies.

| Recipient | Viable cells | | | |
|---|---|---|---|---|
| *P. polymyxa* | 1.2E+07 ± 1.0E+06 | | | |
| **Targeted modifications** | **Targeted position** | **Exconjugants** | **Conjugation efficiency** | **Editing efficiency (%)** |
| *pep* cluster deletion | *pep*-sg1 | 0 | 0 | 0 |
| | *pep*-sg2 | 12 ± 7 | 1.0E-06 ± 5.9E-07 | 91.7 ± 4.2 |
| | *pep*-sg3 | 40 ± 29 | 3.3E-06 ± 2.4E-06 | 100 ± 0 |
| | *pep*-sg4 | 45 ± 18 | 3.8E-06 ± 1.5E-06 | 100 ± 0 |
| | *pep*-sg5 | 19 ± 6 | 1.6E-06 ± 4.8E-07 | 100 ± 0 |
| *dhb* cluster deletion | *dhb*-sg1 | 6 ± 1 | 4.7E-07 ± 7.9E-08 | 100 ± 0 |
| | *dhb*-sg2 | 5 ± 0 | 4.4E-07 ± 3.9E-08 | 100 ± 0 |
| *fus* cluster deletion | *fus*-sg1 | 1 ± 1 | 5.6E-08 ± 7.9E-08 | 0 |
| | *fus*-sg2 | 10 ± 4 | 8.6E-07 ± 3.2E-07 | 96.7 ± 4.7 |
| *pmx* cluster deletion | *pmx*-sg1 | 3 ± 3 | 2.5E-07 ± 2.4E-07 | 0 |
| | *pmx*-sg2 | 6 ± 2 | 4.7E-07 ± 1.4E-07 | 38.9 ± 7.9 |
| *pepC* gene deletion | *pepC* gene | 2.3E+03 ± 1.0E+03 | 1.8E-04 ± 8.5E-05 | 93.9 ± 8.6 |
| *sacB* gene deletion | *sacB* gene | 1.2E+03 ± 7.8E+02 | 9.9E-05 ± 1.4E-06 | 100 ± 0 |
| *pepC-sacB* multiplex gene deletion | *pepC* and *sacB* genes | 17 ± 12 | 1.4E-06 ± 1.0E-06 | 85.0 ± 10.8 |
| *pepC::mRFP-sacB::mRFP* multiplex gene integration | *pepC* and *sacB* genes | 4 ± 1 | 3.1E-07 ± 1.0E-07 | 40.0 ± 20.4 |
| *pepC::sfGFP-sacB::sfGFP* multiplex gene integration | *pepC* and *sacB* genes | 16 ± 1 | 1.3E-06 ± 4,2E-08 | 57.7 ± 11.0 |
| *pep-dhb* multiplex cluster deletion | *pep* and *dhb* clusters | 17 ± 12 | 1.4E-06 ± 1.0E-06 | 82.7 ± 2.7 |

**Table S6**. Biosynthetic gene clusters for antimicrobial compounds in *P. polymyxa* DSM 365*.

| **Cluster** | **Product** | **Antimicrobial type** | **Size (bp)** |
|---|---|---|---|
| *fus* | fusaricidin B | NRPS | 30,709 |
| *pae* | paenicidin B | Lanthipeptide-class-i | 9,122 |
| *pmx* | polymyxin | NRPS | 41,016 |
| *phn* | paenilipoheptin | NRPS | 25,155 |
| *trb* | tridecaptin | NRPS | 58,483 |
| *pnl* | paenilan | Lanthipeptide-class-i | 13,192 |
| *dhb* | bacillibactin | NRPS | 12,417 |
| *paen* | paenibacillin | Lanthipeptide-class-i | 11,575 |

*Analyzed using antiSMASH[4] and manual curation. Only regions with similarity > 50 % to currently known clusters were considered.

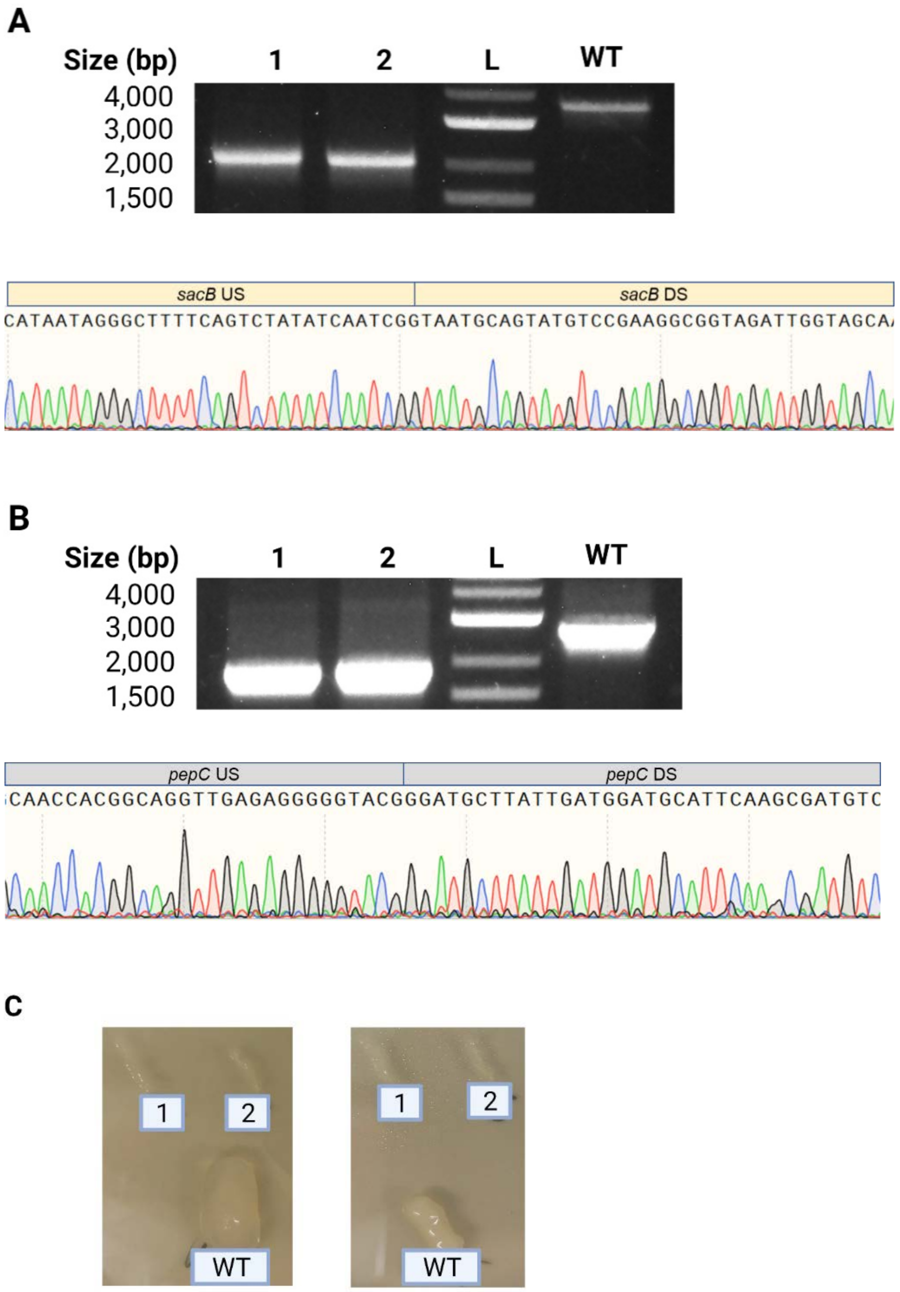

**Figure S1**. Screening of exconjugants of the multiplex gene deletions experiments. PCR screening and sequencing were performed to confirm the deletion of *sacB* (A) and *pepC* (B) genes. Colony PCR was performed on the exconjugants (1-2) and wild type (WT) strains using primers binding outside the homologous regions. (C) Growth of double knockout mutants on EPS-glucose (left) and EPS-sucrose plate (right) with the WT strain as comparison.

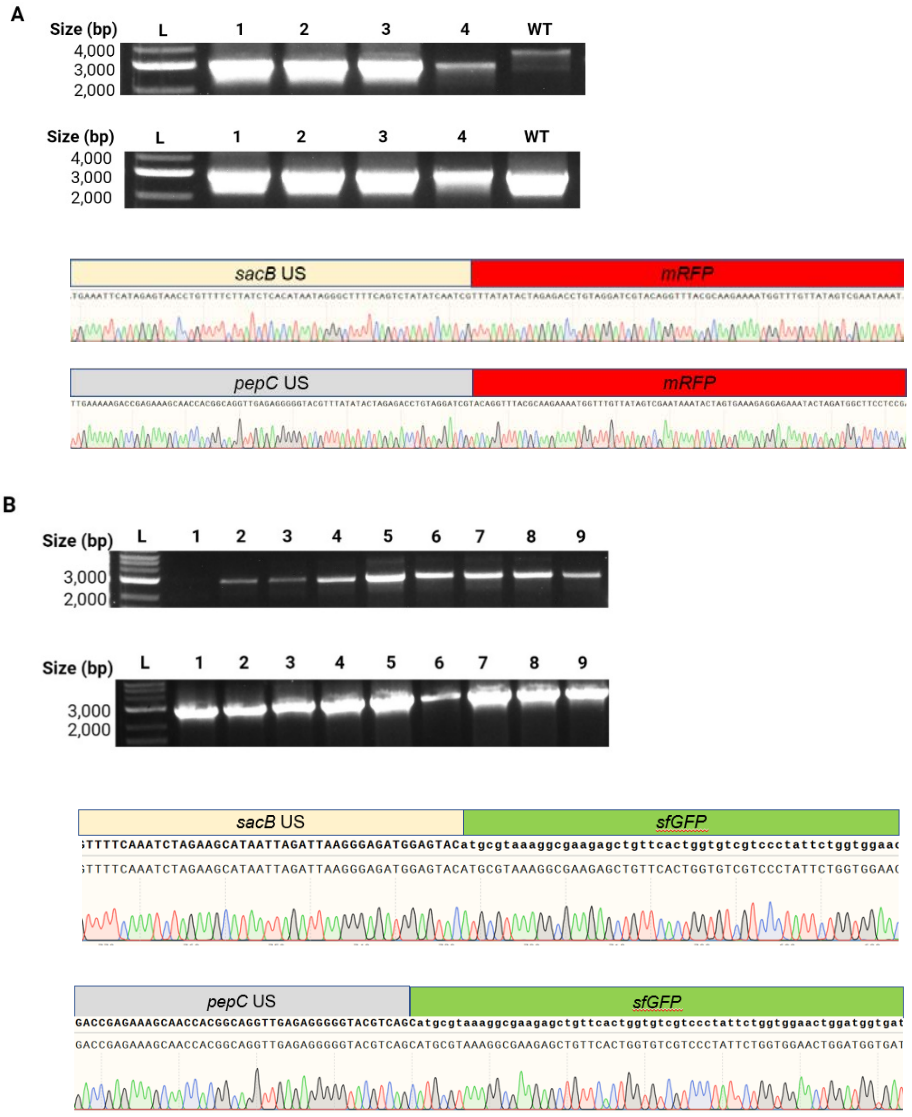

**Figure S2**. Screening of exconjugants of the multiplex gene integrations experiments. (A) PCR screening and sequencing were performed to confirm *mRFP* integrations in the place of *sacB* (top) and *pepC* (bottom) genes. (B) PCR screening and sequencing were performed to confirm *sfGFP* integrations in the place of *sacB* (top) and *pepC* (bottom) genes. Colony PCR was performed on the exconjugants (1-9) and wild type (WT) strains using primers binding outside the homologous regions.

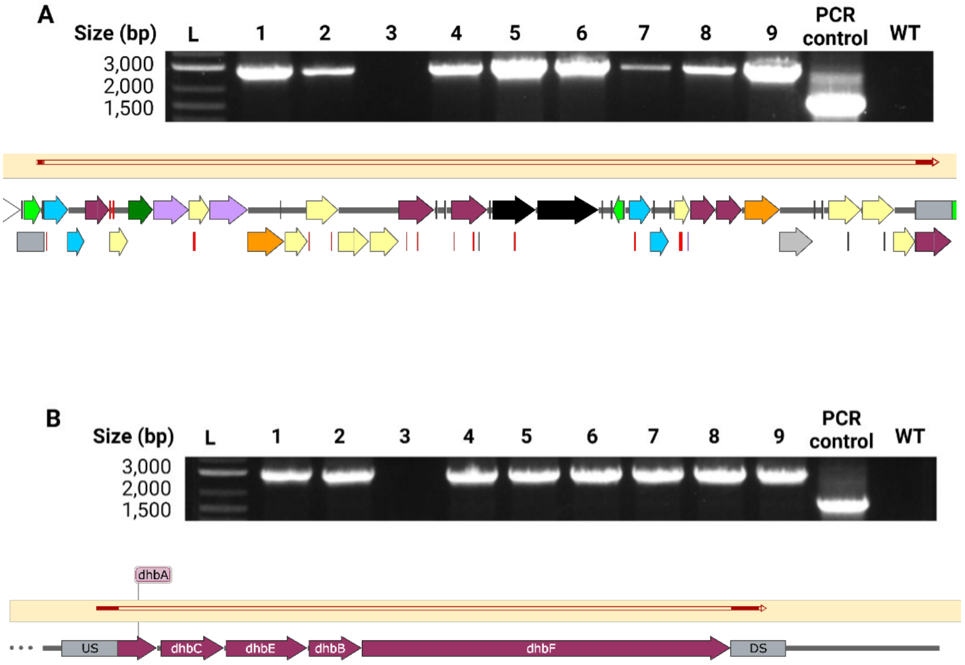

**Figure S3**. Screening of exconjugants of the multiplex cluster deletions experiments. PCR screening and sequencing were performed to confirm *pep* (A) and *dhb* (B) clusters deletion. Colony PCR was performed on the exconjugants (1-9) and wild type (WT) strains using primers binding outside the homologous regions. PCR control was used to ensure viability of the colony PCR by amplification of 16S rRNA.
